# Association between Mitochondrial DNA Copy Number and Sudden Cardiac Death: Findings from the Atherosclerosis Risk in Communities Study (ARIC)

**DOI:** 10.1101/113878

**Authors:** Yiyi Zhang, Eliseo Guallar, Foram N. Ashar, Ryan J. Longchamps, Christina A. Castellani, John Lane, Megan L. Grove, Josef Coresh, Nona Sotoodehnia, Leonard Ilkhanoff, Eric Boerwinkle, Nathan Pankratz, Dan E. Arking

## Abstract

**Aims:** Sudden cardiac death (SCD) is a major public health burden. Mitochondrial dysfunction has been implicated in a wide range of cardiovascular diseases including cardiomyopathy, heart failure, and arrhythmias, but it is unknown if it also contributes to SCD risk. We sought to examine the prospective association between mtDNA copy number (mtDNA-CN), a surrogate marker of mitochondrial function, and SCD risk.

**Methods and Results:** We measured baseline mtDNA-CN in 11,093 participants from the Atherosclerosis Risk in Communities (ARIC) study. mtDNA-CN was calculated from probe intensities of mitochondrial single nucleotide polymorphisms (SNP) on the Affymetrix Genome-Wide Human SNP Array 6.0. SCD was defined as a sudden pulseless condition presumed due to a ventricular tachyarrhythmia in a previously stable individual without evidence of a non-cardiac cause of cardiac arrest. SCD cases were reviewed and adjudicated by an expert committee. During a median follow-up of 20.4 years, we observed 361 SCD cases. After adjusting for age, race, sex, and center, the hazard ratio (HR) for SCD comparing the 1^st^ to the 5^th^ quintiles of mtDNA-CN was 2.24 (95% CI 1.58 to 3.19; p-trend <0.001). When further adjusting for traditional CVD risk factors, prevalent CHD, heart rate, and QT interval duration, the association remained statistically significant. Spline regression models showed that the association was approximately linear over the range of mtDNA-CN values. No apparent interaction by race or by sex was detected.

**Conclusion:** In this community-based prospective study, mtDNA-CN in peripheral blood was inversely associated with the risk of SCD.

## INTRODUCTION

Sudden cardiac death (SCD) is a major clinical and public health burden, with 200,000 to 450,000 events in the United States annually.^1^ Despite major advances in understanding the mechanisms underlying SCD, prediction of SCD remains a challenge, as known clinical factors and biomarkers have only demonstrated limited prognostic power of SCD risk.^2^ Consequently, it is of critical importance to identify novel markers of SCD, particularly among general population samples in which the majority of SCD cases occur.^3^

Mitochondria are cellular organelles specialized in energy production and responsible for generating nearly all cellular adenosine triphosphate (ATP) through oxidative phosphorylation.^4^ Mitochondrial dysfunction may contribute to a wide range of conditions including cardiovascular diseases (CVD) such as cardiomyopathy, heart failure, and arrhythmias.^5-7^ Each cell has multiple mitochondria, and each mitochondrion has 2 to 10 copies of mtDNA.^8^ Mitochondrial DNA copy number (mtDNA-CN), which measures the amount of mtDNA relative to the amount of nuclear DNA, has been established as an indirect marker of mitochondrial function.^9^ Furthermore, mtDNA-CN is measured using a low-cost scalable assay and allows for rapid determination of mitochondrial function in a large number of samples. Indeed, recent studies have shown that low levels of mtDNA-CN were associated with diabetes, chronic kidney disease, CVD, and mortality in general population samples.^10-14^ However, it is unknown if mtDNA-CN is also a risk marker of SCD.

In the present study, we sought to examine the prospective association between baseline mtDNA-CN and the risk of SCD among participants from the Atherosclerosis Risk in Communities (ARIC) study. We hypothesized that participants with lower levels of mtDNA-CN would be at increased risk of SCD.

## METHODS

### Study Design and Population

The ARIC study is a population-based, prospective cohort study of 15,792 individuals 45 to 64 years of age from 4 US communities (Forsyth County, NC; Jackson, MS; suburban Minneapolis, MN; and Washington County, MD).^15^ The first visit was carried out during 1987–1989, with 4 subsequent in-person visits and annual telephone interviews (a fifth follow-up visit is currently underway). The present analysis was restricted to 11,455 Caucasian or African-American participants who had DNA available for mtDNA-CN measurements. We further excluded 362 participants with missing CVD risk factor information or other covariates. The final sample size was 11,093 (4,971 men and 6,122 women). All centers obtained approval from their respective institutional review boards and all participants provided written informed consent.

### Clinical Data Collection

ARIC participants underwent a comprehensive medical history and cardiovascular examination by centrally trained clinical teams during each standardized clinical visit.^15^ Age, sex, race/ethnicity, and smoking status were self-reported. Height and weight were measured and body mass index was calculated. Hypertension was defined as systolic blood pressure ≥ 140 mm Hg, diastolic blood pressure ≥ 90 mm Hg, or current use of anti-hypertension medication.

Fasting blood samples were drawn and sent to a central laboratory for measurement of glucose, lipids, and creatinine. Diabetes was defined as fasting glucose ≥ 126 mg/dL, non-fasting glucose ≥ 200 mg/dL, use of hypoglycemic medication, or self-reported physician diagnosis of diabetes. Estimated Glomerular filtration rate (eGFR) was calculated using the Chronic Kidney Disease Epidemiology Collaboration (CKD-EPI) equation.^16^ Prevalent coronary heart disease (CHD) was defined as a history of definite or probable myocardial infarction or cardiac procedure. Heart rate and QT interval duration was measured by standard 12-lead electrocardiogram (ECG).

### Sudden Cardiac Death Events

SCD was defined as a sudden pulseless condition presumed due to a ventricular tachyarrhythmia in a previously stable individual without evidence of a non-cardiac cause of cardiac arrest. To identify SCD, cases of fatal cardiovascular death that occurred by December 31, 2012 were reviewed and adjudicated by a committee of electrophysiologists, general cardiologists, and internists in two phases. In the first phase, CVD deaths occurring on or before December 31, 2001 were adjudicated by 5 physicians. In the second phase, CVD deaths occurring between January 1, 2002 and December 31, 2012 were adjudicated by a committee of 11 physicians. All cases of fatal CVD that occurred out of the hospital or in the emergency room were reviewed. In the first phase, all in-hospital CVD deaths were also reviewed. In the second phase, in-hospital deaths were reviewed only if cardiac arrest with cardiopulmonary resuscitation (CPR) occurred prior to hospital admission.

The committee reviewed all available data from death certificates, informant interviews, physician questionnaires, coroner reports, prior medical history, and hospital discharge summaries, in addition to circumstances surrounding the event, to help classify whether the subject had experienced SCD. For witnessed cases, SCD was operationally defined as a sudden pulseless condition without evidence of a non-cardiac cause of cardiac arrest. For unwitnessed cases, the patient had to have been known to be in a stable condition in the 24 hours prior to cardiac arrest without evidence of a non-cardiac cause of the cardiac arrest. Study participants were not classified as sudden death if there was evidence of an acute non-cardiac morbidity that could account for the arrest, such as drug overdose, stroke, aortic aneurysm rupture, other acute bleeding, pulmonary embolism, acute respiratory failure, or trauma. To minimize misclassification of phenotype, we excluded cases with chronic terminal illnesses, including terminal cancer and end-stage liver disease. Severe dementia or severely debilitated individuals who required long-term care were excluded. Similarly, nursing home deaths were excluded. Inter-reviewer agreement for SCD adjudication was 83.2%, and there was 92.5% agreement between the two phases of SCD adjudication.

### Measurement of mtDNA Copy Number

The method for measuring mtDNA-CN has been described previously.^13^ Briefly, DNA samples derived from buffy coat were used for array-based genotyping. mtDNA-CN was calculated from probe intensities of mitochondrial single nucleotide polymorphisms (SNP) on the Affymetrix Genome-Wide Human SNP Array 6.0 using the Genvisis software package (http://www.genvisis.org). This method uses median mitochondrial probe intensity of 25 high-quality mitochondrial probes as an initial raw measure of mtDNA-CN. To correct for batch effects, DNA quality, and starting DNA quantity, surrogate variable analysis was applied to probe intensities of 43,316 autosomal SNPs to adjust for different amounts of total DNA hybridized to the arrays and additional technical artifacts.^17^ We used linear regression model to adjust the effect of age, sex, enrollment center, and the surrogate variables on the initial raw estimate of mtDNA-CN. In the present analysis, the standardized residuals from this linear model were used as the measurement for mtDNA-CN (i.e., with a mean of 0 and standard deviation of 1).

### Statistical Analyses

DNA for mtDNA-CN analysis was collected in visit 1 (1987-1989) for 470 participants (4.2%), in visit 2 (1990-1992) for 8,810 participants (79.4%), in visit 3 (1993-1995) for 1,752 participants (15.8%), and in visit 4 for (1996-1998) for 61 participants (0.6%). The visit of DNA collection in each participant was considered the baseline visit and all covariates were obtained from that visit. Follow-up for events started from the baseline visit and continued until death or through December 31, 2012, whichever came first.

Baseline characteristics of the study population were shown by quintiles of mtDNA-CN. We used one-way analysis-of-variance (ANOVA) to compare means for continuous variables, and *χ*^2^ tests to compare proportions for categorical variables. We used a Cox proportional hazards model to estimate hazard ratios (HR) and 95% confidence intervals (CI) for the association between mtDNA-CN and SCD. In the primary analysis, we categorized mtDNA-CN into quintiles based on the sample distribution. Tests for linear trend across quintiles were conducted by including a variable with the median mtDNA-CN level of each quintile in the models. In secondary analysis, we modeled mtDNA as a continuous variable and estimated the HR for SCD comparing the 10^th^ to the 90^th^ percentile of mtDNA-CN. Additionally, we modeled mtDNA-CN as restricted quadratic splines with knots at the 5^th^, 50^th^, and 95^th^ percentiles of its distribution to provide a smooth yet flexible description of the dose-response relationship between mtDNA-CN and SCD.

To adjust for potential confounders, we used 4 multivariate models with progressive degrees of adjustment. The first model was adjusted for age, race, sex, and enrollment center. The second model was further adjusted for body mass index, smoking, total and HDL cholesterol, triglycerides, hypertension, and diabetes. The third model was further adjusted for heart rate, and QT interval. The fourth model was further adjusted for prevalent CHD at baseline. The proportional hazards assumption was checked by plotting the log(-log(survival)) vs. log(survival time) and by using Schoenfeld Residuals.

Additionally, we performed pre-specified subgroup analyses by race and sex, and tested for potential interactions. We also performed several sensitivity analyses. First, we excluded 802 participants with prevalent CHD at baseline. Second, since mtDNA-CN from peripheral blood is associated with white blood cell count (WBC), we further adjusted for log-transformed WBC count among 9,435 participants in whom WBC measurements were available. Third, we adjusted for eGFR among 9,341 participants in whom creatinine measurements were available. Finally, we performed competing risk analyses treating deaths other than SCD as competing events, with similar results (data not shown). All statistical analyses were performed using STATA version 12 (StataCorp LP, College Station, Texas).

## RESULTS

The average age (SD) of study participants at baseline was 57.9 (6.0) years (Table 1); 44.8% of participants were men and 79.4% were white. Participants with lower mtDNA-CN were more likely to be current smoker, hypertensive, diabetic, and to have lower HDL cholesterol, higher triglycerides, faster heart rate, slightly shorter QT interval, and prevalent CHD at baseline. As expected, no association with age or sex was observed as our mtDNA-CN measure was adjusted for age, sex, and enrollment center.

**Table 1.**
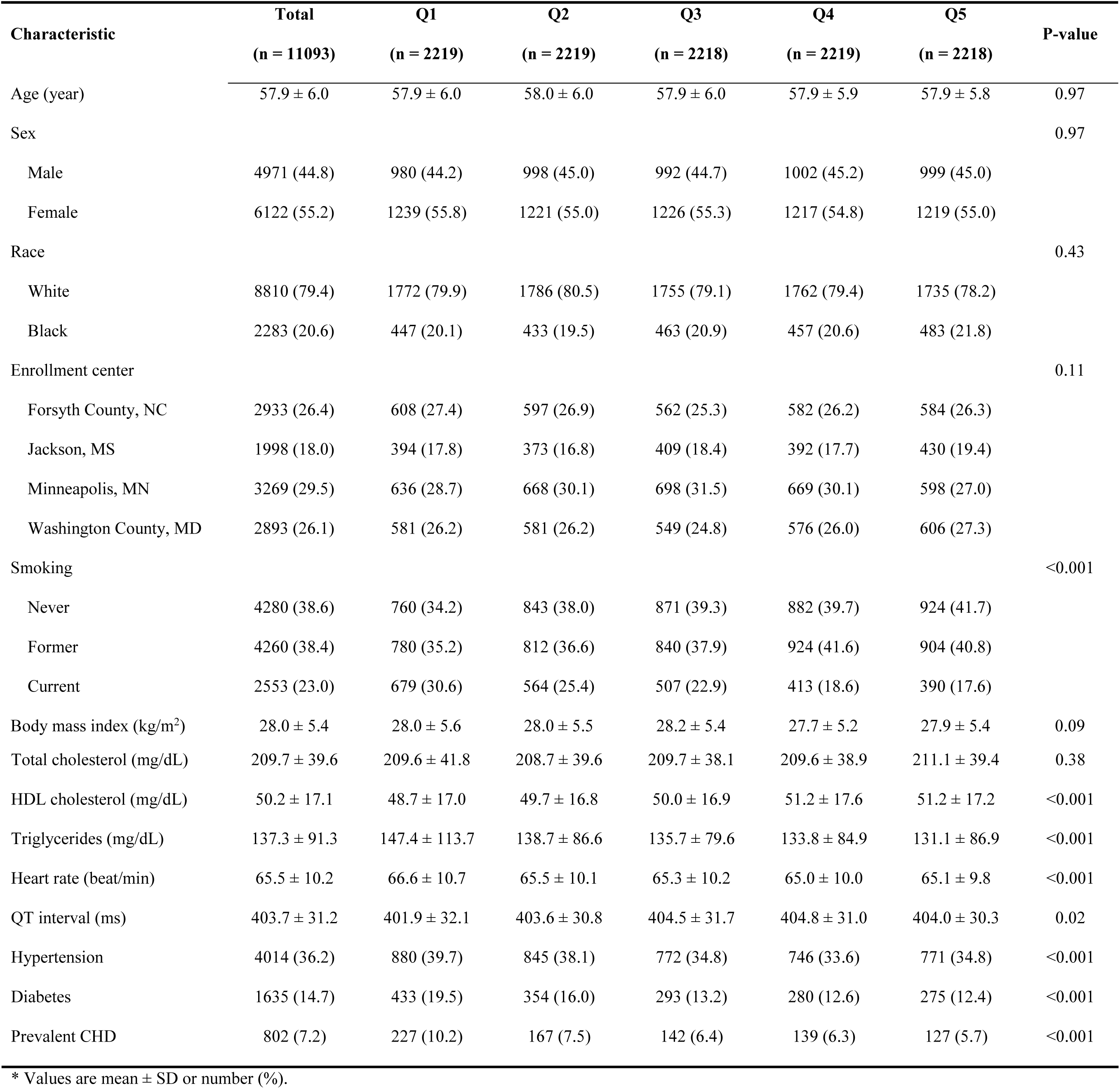
Baseline characteristics of study participants by quintiles of mtDNA copy number. *.

| **Characteristic** | **Total**<br>**(n = 11093)** | **Q1**<br>**(n = 2219)** | **Q2**<br>**(n = 2219)** | **Q3**<br>**(n = 2218)** | **Q4**<br>**(n = 2219)** | **Q5**<br>**(n = 2218)** | **P-value** |
| --- | --- | --- | --- | --- | --- | --- | --- |
| Age (year) | 57.9 ± 6.0 | 57.9 ± 6.0 | 58.0 ± 6.0 | 57.9 ± 6.0 | 57.9 ± 5.9 | 57.9 ± 5.8 | 0.97 |
| Sex |  |  |  |  |  |  | 0.97 |
| Male | 4971 (44.8) | 980 (44.2) | 998 (45.0) | 992 (44.7) | 1002 (45.2) | 999 (45.0) |  |
| Female | 6122 (55.2) | 1239 (55.8) | 1221 (55.0) | 1226 (55.3) | 1217 (54.8) | 1219 (55.0) |  |
| Race |  |  |  |  |  |  | 0.43 |
| White | 8810 (79.4) | 1772 (79.9) | 1786 (80.5) | 1755 (79.1) | 1762 (79.4) | 1735 (78.2) |  |
| Black | 2283 (20.6) | 447 (20.1) | 433 (19.5) | 463 (20.9) | 457 (20.6) | 483 (21.8) |  |
| Enrollment center |  |  |  |  |  |  | 0.11 |
| Forsyth County, NC | 2933 (26.4) | 608 (27.4) | 597 (26.9) | 562 (25.3) | 582 (26.2) | 584 (26.3) |  |
| Jackson, MS | 1998 (18.0) | 394 (17.8) | 373 (16.8) | 409 (18.4) | 392 (17.7) | 430 (19.4) |  |
| Minneapolis, MN | 3269 (29.5) | 636 (28.7) | 668 (30.1) | 698 (31.5) | 669 (30.1) | 598 (27.0) |  |
| Washington County, MD | 2893 (26.1) | 581 (26.2) | 581 (26.2) | 549 (24.8) | 576 (26.0) | 606 (27.3) |  |
| Smoking |  |  |  |  |  |  | <0.001 |
| Never | 4280 (38.6) | 760 (34.2) | 843 (38.0) | 871 (39.3) | 882 (39.7) | 924 (41.7) |  |
| Former | 4260 (38.4) | 780 (35.2) | 812 (36.6) | 840 (37.9) | 924 (41.6) | 904 (40.8) |  |
| Current | 2553 (23.0) | 679 (30.6) | 564 (25.4) | 507 (22.9) | 413 (18.6) | 390 (17.6) |  |
| Body mass index (kg/m<sup>2</sup>) | 28.0 ± 5.4 | 28.0 ± 5.6 | 28.0 ± 5.5 | 28.2 ± 5.4 | 27.7 ± 5.2 | 27.9 ± 5.4 | 0.09 |
| Total cholesterol (mg/dL) | 209.7 ± 39.6 | 209.6 ± 41.8 | 208.7 ± 39.6 | 209.7 ± 38.1 | 209.6 ± 38.9 | 211.1 ± 39.4 | 0.38 |
| HDL cholesterol (mg/dL) | 50.2 ± 17.1 | 48.7 ± 17.0 | 49.7 ± 16.8 | 50.0 ± 16.9 | 51.2 ± 17.6 | 51.2 ± 17.2 | <0.001 |
| Triglycerides (mg/dL) | 137.3 ± 91.3 | 147.4 ± 113.7 | 138.7 ± 86.6 | 135.7 ± 79.6 | 133.8 ± 84.9 | 131.1 ± 86.9 | <0.001 |
| Heart rate (beat/min) | 65.5 ± 10.2 | 66.6 ± 10.7 | 65.5 ± 10.1 | 65.3 ± 10.2 | 65.0 ± 10.0 | 65.1 ± 9.8 | <0.001 |
| QT interval (ms) | 403.7 ± 31.2 | 401.9 ± 32.1 | 403.6 ± 30.8 | 404.5 ± 31.7 | 404.8 ± 31.0 | 404.0 ± 30.3 | 0.02 |
| Hypertension | 4014 (36.2) | 880 (39.7) | 845 (38.1) | 772 (34.8) | 746 (33.6) | 771 (34.8) | <0.001 |
| Diabetes | 1635 (14.7) | 433 (19.5) | 354 (16.0) | 293 (13.2) | 280 (12.6) | 275 (12.4) | <0.001 |
| Prevalent CHD | 802 (7.2) | 227 (10.2) | 167 (7.5) | 142 (6.4) | 139 (6.3) | 127 (5.7) | <0.001 |

\* Values are mean ± SD or number (%).

During a median follow-up of 20.4 years, 361 participants had a SCD event (Table 2). In a model adjusted for age, race, sex, and enrollment center, the HR for SCD comparing the 1^st^ quintile of mtDNA-CN to the 5^th^ quintile was 2.24 (95% CI 1.58 to 3.19; *p*-trend <0.001). When further adjusted for additional CVD risk factors, heart rate, QT interval, and prevalent CHD, the HR was attenuated but remained statistically significant (HR 1.74, 95% CI 1.21 to 2.50; *p*-trend = 0.003). Further adjustment for WBC (HR = 1.65, 95% CI 1.10 to 2.46; *p*-trend = 0.01) or eGFR (HR = 1.65, 95% CI 1.11 to 2.46; *p*-trend = 0.01) did not appreciably change the magnitude of the association. Spline regression analysis also confirmed that mtDNA-CN was inversely associated with the risk of SCD, with an approximately linear dose-response relationship (p-value for non-linear spline terms > 0.85; Figure 1).

**Table 2.**
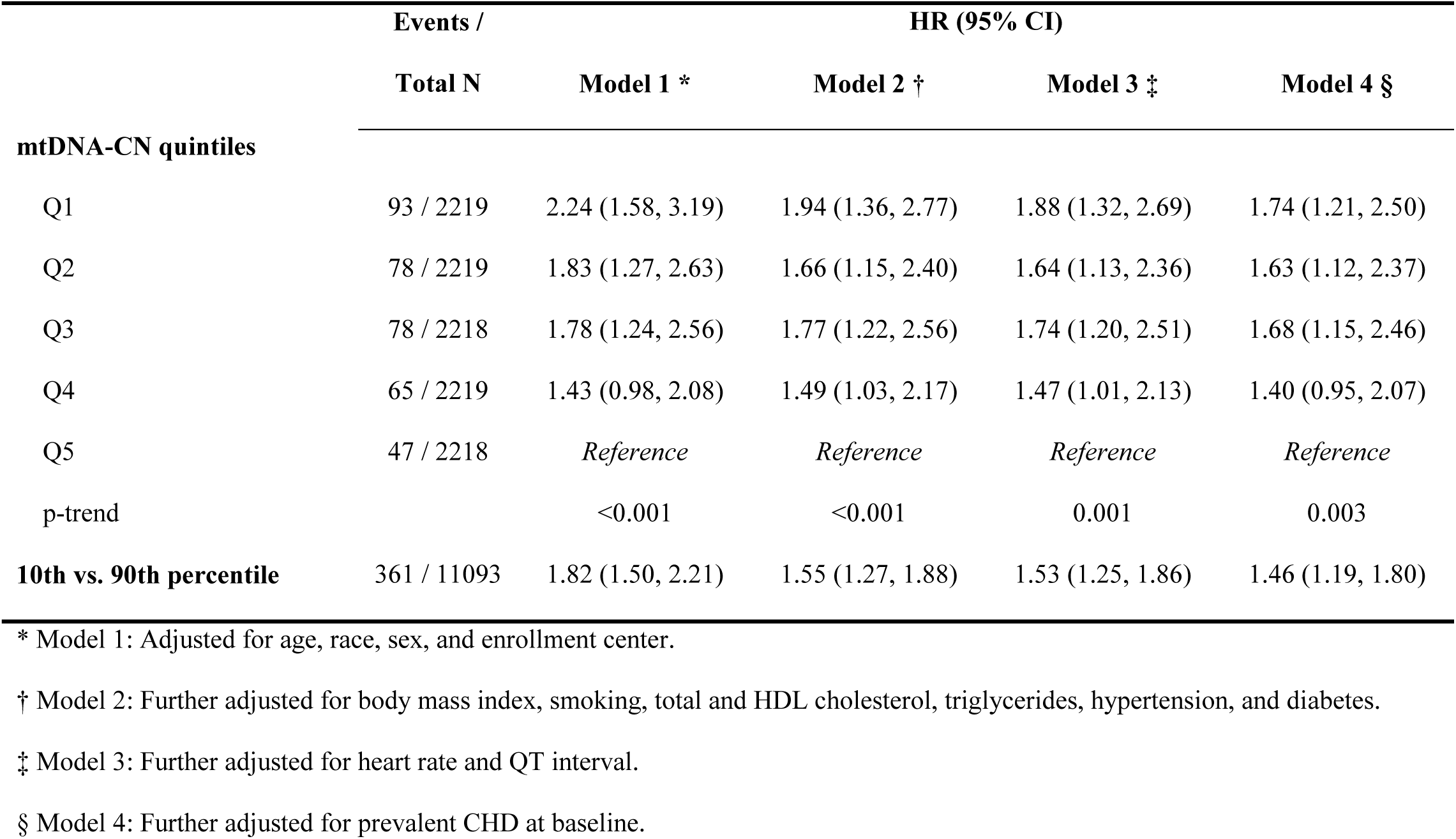
Hazard ratio for sudden cardiac death (SCD) by quintiles of mtDNA copy number.

**Figure 1.**
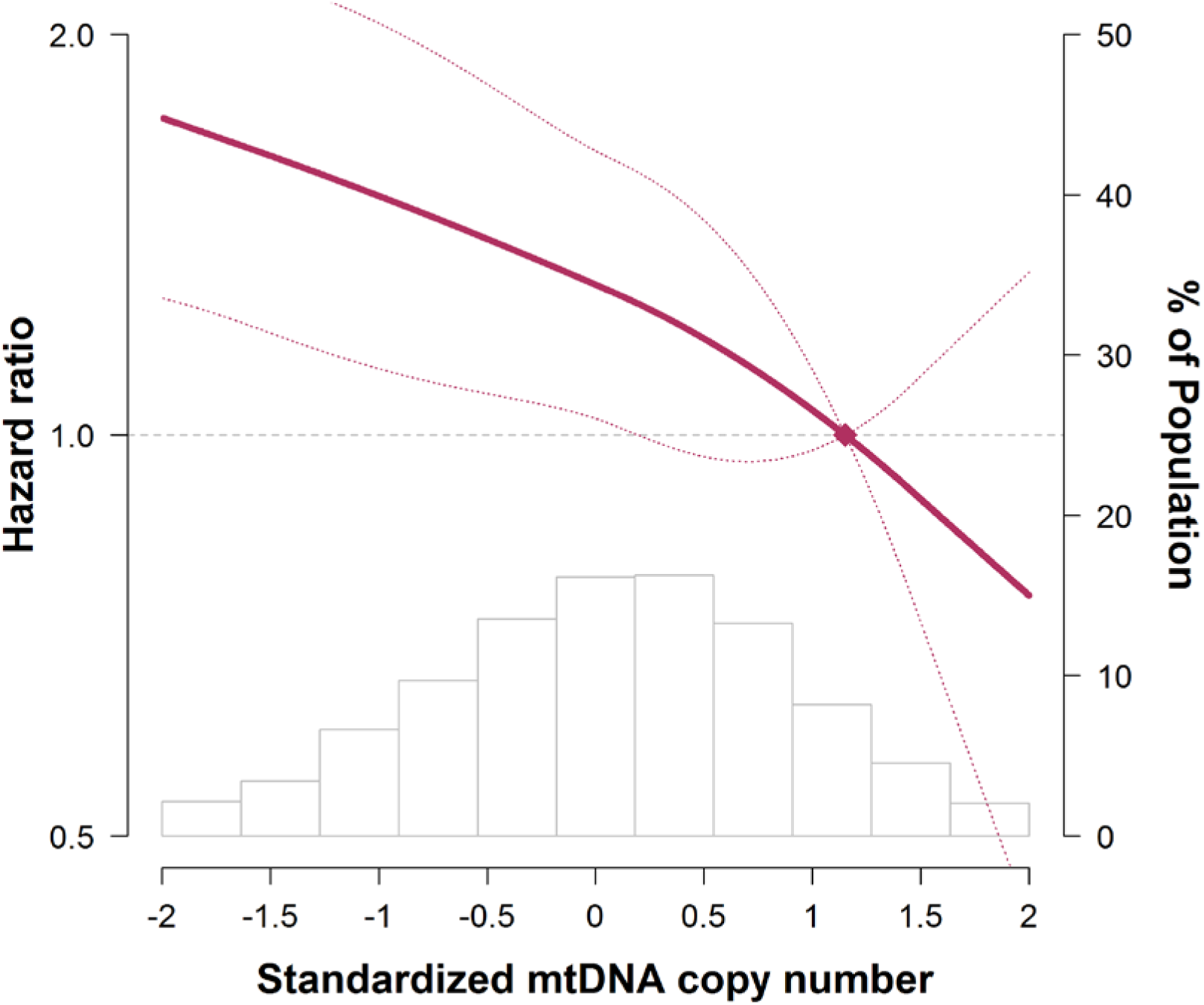
Hazard ratios for sudden cardiac death by levels of mtDNA copy number. The curves represent adjusted hazard ratios (solid line) and their 95% confidence intervals (dashed lines) based on restricted quadratic splines of mtDNA copy number with knots at the 5^th^, 50^th^, and 95^th^ percentiles of its distribution. The reference value (diamond dot) was set at the 90^th^ percentile of the distribution. Results were obtained from a Cox model adjusted for age, race, sex, enrollment center, body mass index, smoking, total and HDL cholesterol, triglycerides, hypertension, diabetes, heart rate, QT interval, and prevalent CHD at baseline. Histograms represent the frequency distribution of mtDNA copy number at baseline.

When stratified by race and sex, the associations between mtDNA-CN and SCD was similar among men and women, and slightly stronger in whites than in blacks, but the interactions by race or by sex were not statistically significant (all *p*-interactions >0.1; Supplemental Tables 1 & 2). In sensitivity analysis, the association between mtDNA-CN and SCD persisted after excluding participants with prevalent CHD at baseline (**Supplemental** Table 3).

## DISCUSSION

In this large community-based prospective cohort of white and black Americans, we found that mtDNA-CN was inversely associated with the risk of SCD. This novel association was approximately linear and similar among men and women, as well as among whites and blacks. Additionally, the association between mtDNA-CN and SCD remained after adjusting for traditional CVD risk factors and prevalent CHD, suggesting that mtDNA-CN may be a risk marker for SCD independent of CHD, traditional CHD risk factors, heart rate, and QT interval duration.

mtDNA dysfunction has been associated with several CVD outcomes including heart failure, cardiomyopathy, long QT syndrome, and arrhythmias.^5, 6 18-20^ However, most of the evidence was obtained in small studies of patients with genetically confirmed mtDNA abnormalities (i.e., mtDNA deletions and/or mutations), and the clinical implication of variations in mtDNA function in the general population is largely unknown. mtDNA-CN, which measures the average level of mtDNA per cell, correlates with mitochondrial enzyme activities and ATP production,^21^ and is a surrogate marker of mtDNA function.^9, 11, 13^ The cost for measuring mtDNA-CN in a typical research setting is <$2 per sample using qPCR. mtDNA-CN could represent a low-cost approach to improving risk prediction of SCD in the general population. Indeed, recent studies in general population samples have shown associations of mtDNA-CN with aging, frailty, chronic kidney disease, and all-cause mortality.^8, 11, 13^

The precise mechanism underlying the association between mtDNA-CN and SCD is unclear. Cardiac cells rely heavily on mitochondrial oxidative energy to maintain myocardial contractility and electrical activity.^7^ It is estimated that 30% of the cardiac ATP generated by mitochondria is used for sarcolemmal and sarcoplasmic reticulum ion channels and transporters, which are required for the electrical activity of the myocytes.^7, 22^ It is possible that decreased mtDNA-CN measured in blood reflects mitochondrial dysfunction in cardiac cells, which could compromise ATP production and energy supply to ion channels and transporters, leading to altered ion homeostasis, membrane excitability, and cardiac arrhythmias.^7^

In addition to ATP, mitochondria generate reactive oxygen species (ROS) as a by-product of oxidative phosphorylation.^7^ Excessive ROS production can alter action potential propagation and impair cardiac excitability through a number of ion channels and transporters.^7, 23, 24^ Furthermore, an increase in ROS above a threshold level triggers the opening of mitochondrial channels such as the mitochondrial permeability transition pore (MPTP) or the inner membrane anion channel (IMAC), leading to the collapse of mitochondrial membrane potential and a further increase in ROS generation, a process known as mitochondrial ROS-induced ROS release.^25-27^ mtDNA is highly susceptible to oxidative damage due to its close proximity to mitochondrial ROS, lack of protective histone proteins, and limited DNA repair capabilities. As a consequence, oxidative stress-related mitochondrial damage may further contribute to a decreased mtDNA-CN and an increased risk of SCD.^8, 28^ Additionally, regional mitochondrial depolarization triggered by oxidative stress activates sarcolemmal ATP-sensitive K^+^ currents to form a metabolic sink, contributing to the development of re-entry arrhythmias.^29^ Mitochondrial oxidative stress also plays an important role in angiotensin II-induced gap junction remodeling and arrhythmogenesis.^30^

A few limitations of this study need to be considered. Due to the observational nature of our study, we could identify an association but not establish a causal link between mtDNA-CN and SCD. Our study measured mtDNA-CN in DNA derived from peripheral blood, which may not necessarily be the relevant tissue with respect to SCD. However, mtDNA-CN in peripheral blood has been shown to correlate with that of cardiomyocytes (Pearson correlation = 0.72),^10^ suggesting that mtDNA-CN in peripheral blood may serve as a marker for mitochondrial function in the heart. Additionally, we measured mtDNA-CN at a single time point and did not account for the dynamic nature of mtDNA-CN throughout the life course. Serial measures of mtDNA-CN may provide additional insight into the relationship between mitochondrial function and SCD.

The major strengths of this study include a large sample size of both white and black participants, prospective cohort study design and long follow-up period for SCD events, stringent SCD adjudication, as well as the use of state-of-the-art tools for mtDNA-CN measurement.

In conclusion, in this community-based prospective study, mtDNA-CN in peripheral blood was inversely associated with SCD risk independent of CHD, CHD risk factors, heart rate, and QT interval duration. These findings may provide a novel risk marker for SCD among general population samples and uncover new therapeutic targets for reducing SCD. Future studies are needed to confirm our findings in other study populations and to elucidate underlying mechanisms.

## ACKNOWLEDGEMENT

We would like to acknowledge the sudden cardiac death mortality classification committee members: Nona Sotoodehnia (lead), Selcuk Adabag, Sunil Agarwal, Lin Chen, Raj Deo, Leonard Ilkhanoff, Liviu Klein, Saman Nazarian, Ashleigh Owen, Kris Patton, and Larisa Tereschchenko.

## FUNGDING SUPPORT

This work was supported by US National Institutes of Health grants R01HL131573 (YZ, EG, FNA, RJL, CAC, DEA), R01HL111267 (FNA, RJL, DEA), and R01HL116747 (NS). The Atherosclerosis Risk in Communities Study is carried out as a collaborative study supported by National Heart, Lung, and Blood Institute contracts (HHSN268201100005C, HHSN268201100006C, HHSN268201100007C, HHSN268201100008C, HHSN268201100009C, HHSN268201100010C, HHSN268201100011C, and HHSN268201100012C), R01HL087641, R01HL59367 and R01HL086694; National Human Genome Research Institute contract U01HG004402; and National Institutes of Health contract HHSN268200625226C. Adjudication of sudden cardiac death cases was funded by R01 HL111089. The authors thank the staff and participants of the ARIC study for their important contributions. Infrastructure was partly supported by Grant Number UL1RR025005, a component of the National Institutes of Health and NIH Roadmap for Medical Research.

## SUPPLEMENTAL MATERIAL

**Supplemental Table 1.** Hazard ratio for sudden cardiac death (SCD) by quintiles of mtDNA copy number, stratified by race.

|  | Events /<br>Total N | HR (95% CI) |  |  |  |
| --- | --- | --- | --- | --- | --- |
|  |  | Model 1 * | Model 2 † | Model 3 ‡ | Model 4 § |
| White |  |  |  |  |  |
| mtDNA-CN quintiles |  |  |  |  |  |
| Q1 | 72 / 1762 | 2.43 (1.61, 3.66) | 1.94 (1.28, 2.93) | 1.86 (1.22, 2.82) | 1.69 (1.11, 2.59) |
| Q2 | 44 / 1762 | 1.43 (0.91, 2.24) | 1.23 (0.78, 1.94) | 1.24 (0.79, 1.96) | 1.21 (0.76, 1.91) |
| Q3 | 50 / 1762 | 1.61 (1.04, 2.51) | 1.51 (0.97, 2.34) | 1.47 (0.94, 2.30) | 1.34 (0.85, 2.12) |
| Q4 | 38 / 1762 | 1.15 (0.72, 1.83) | 1.18 (0.74, 1.87) | 1.17 (0.74, 1.87) | 1.15 (0.71, 1.84) |
| Q5 | 33 / 1762 | Reference | Reference | Reference | Reference |
| p-trend |  | <0.001 | 0.002 | 0.004 | 0.01 |
| 10th vs. 90th percentile | 237 / 8810 | 2.06 (1.65, 2.58) | 1.69 (1.33, 2.14) | 1.68 (1.32, 2.14) | 1.58 (1.22, 2.04) |
| Black |  |  |  |  |  |
| mtDNA-CN quintiles |  |  |  |  |  |
| Q1 | 21 / 457 | 1.81 (0.90, 3.63) | 1.85 (0.91, 3.78) | 1.85 (0.91, 3.76) | 1.78 (0.87, 3.65) |
| Q2 | 36 / 457 | 3.04 (1.61, 5.73) | 3.08 (1.62, 5.86) | 3.01 (1.58, 5.74) | 3.12 (1.61, 6.05) |
| Q3 | 28 / 456 | 2.19 (1.13, 4.24) | 2.54 (1.30, 4.97) | 2.57 (1.31, 5.04) | 2.67 (1.34, 5.33) |
| Q4 | 26 / 457 | 2.11 (1.08, 4.12) | 2.25 (1.15, 4.39) | 2.19 (1.11, 4.32) | 2.10 (1.05, 4.24) |
| Q5 | 13 / 456 | Reference | Reference | Reference | Reference |
| p-trend |  | 0.04 | 0.05 | 0.05 | 0.06 |
| **10th vs. 90th percentile** | 124 / 2283 | 1.31 (0.95, 1.81) | 1.26 (0.92, 1.72) | 1.24 (0.91, 1.70) | 1.24 (0.90, 1.70) |

P-interaction by race was 0.02 in Model 1; all other p-interactions by race were > 0.1.
\* Model 1: Adjusted for age, sex, and enrollment center.
† Model 2: Further adjusted for body mass index, smoking, total and HDL cholesterol, triglycerides, hypertension, and diabetes.
‡ Model 3: Further adjusted for heart rate and QT interval.
§ Model 4: Further adjusted for prevalent CHD at baseline.

**Supplemental Table 2.** Hazard ratio for sudden cardiac death (SCD) by quintiles of mtDNA copy number, stratified by sex.

|  | Events /<br>Total N | HR (95% CI) |  |  |  |
| --- | --- | --- | --- | --- | --- |
|  |  | Model 1 * | Model 2 † | Model 3 ‡ | Model 4 § |
|  |  | Male |  |  |  |
| mtDNA-CN quintiles |  |  |  |  |  |
| Q1 | 61 / 995 | 2.49 (1.58, 3.91) | 2.17 (1.38, 3.41) | 2.09 (1.32, 3.30) | 1.95 (1.22, 3.11) |
| Q2 | 44 / 994 | 1.77 (1.10, 2.86) | 1.56 (0.97, 2.53) | 1.54 (0.95, 2.49) | 1.57 (0.96, 2.58) |
| Q3 | 53 / 994 | 2.10 (1.32, 3.34) | 2.04 (1.28, 3.26) | 1.99 (1.25, 3.19) | 1.88 (1.16, 3.06) |
| Q4 | 47 / 994 | 1.74 (1.09, 2.79) | 1.82 (1.13, 2.93) | 1.80 (1.12, 2.89) | 1.76 (1.08, 2.87) |
| Q5 | 27 / 994 | Reference | Reference | Reference | Reference |
| p-trend |  | <0.001 | 0.004 | 0.01 | 0.02 |
| 10th vs. 90th percentile | 232 / 4971 | 1.91 (1.51, 2.41) | 1.65 (1.30, 2.11) | 1.64 (1.28, 2.10) | 1.56 (1.20, 2.02) |
|  |  | Female |  |  |  |
| mtDNA-CN quintiles |  |  |  |  |  |
| Q1 | 33 / 1225 | 1.98 (1.14, 3.44) | 1.58 (0.89, 2.80) | 1.61 (0.90, 2.88) | 1.51 (0.85, 2.70) |
| Q2 | 33 / 1224 | 1.90 (1.09, 3.31) | 1.79 (1.02, 3.16) | 1.81 (1.02, 3.23) | 1.76 (0.99, 3.12) |
| Q3 | 27 / 1225 | 1.45 (0.81, 2.57) | 1.46 (0.81, 2.63) | 1.49 (0.83, 2.70) | 1.52 (0.84, 2.76) |
| Q4 | 16 / 1224 | 0.88 (0.46, 1.69) | 0.86 (0.45, 1.63) | 0.82 (0.43, 1.58) | 0.76 (0.38, 1.50) |
| Q5 | 20 / 1224 | 1.00 (1.00, 1.00) | 1.00 (1.00, 1.00) | 1.00 (1.00, 1.00) | 1.00 (1.00, 1.00) |
| p-trend |  | 0.001 | 0.02 | 0.01 | 0.02 |
| 10th vs. 90th percentile | 129 / 6122 | 1.67 (1.21, 2.32) | 1.34 (0.97, 1.84) | 1.35 (0.99, 1.85) | 1.32 (0.95, 1.83) |

All p-interactions by sex were $> 0.1$ .
\* Model 1: Adjusted for age, race, and enrollment center.
† Model 2: Further adjusted for body mass index, smoking, total and HDL cholesterol, triglycerides, hypertension, and diabetes.
‡ Model 3: Further adjusted for heart rate and QT interval.
§ Model 4: Further adjusted for prevalent CHD at baseline.

**Supplemental Table 3.** Hazard ratio for sudden cardiac death (SCD) by quintiles of mtDNA copy number, excluding participants with prevalent CHD at baseline.

|  | Events /<br>Total N | HR (95% CI) |  |  |
| --- | --- | --- | --- | --- |
|  |  | Model 1 * | Model 2 † | Model 3 ‡ |
| mtDNA-CN quintiles |  |  |  |  |
| Q1 | 58 / 2059 | 1.69 (1.12, 2.54) | 1.51 (1.00, 2.28) | 1.47 (0.97, 2.21) |
| Q2 | 52 / 2058 | 1.48 (0.98, 2.25) | 1.36 (0.89, 2.07) | 1.35 (0.89, 2.05) |
| Q3 | 58 / 2058 | 1.59 (1.06, 2.39) | 1.65 (1.09, 2.49) | 1.64 (1.09, 2.48) |
| Q4 | 48 / 2058 | 1.30 (0.85, 1.98) | 1.40 (0.92, 2.13) | 1.40 (0.92, 2.14) |
| Q5 | 39 / 2058 | Reference | Reference | Reference |
| p-trend |  | 0.01 | 0.08 | 0.12 |
| 10th vs. 90th percentile | 255 /<br>10291 | 1.63 (1.26, 2.10) | 1.40 (1.09, 1.80) | 1.36 (1.06, 1.75) |

\* Model 1: Adjusted for age, race, sex, and enrollment center.
† Model 2: Further adjusted for body mass index, smoking, total and HDL cholesterol, triglycerides, hypertension, and diabetes.
‡ Model 3: Further adjusted for heart rate and QT interval.

## ABBREVIATIONS

ANOVA: analysis-of-variance
ARIC: Atherosclerosis Risk in Communities
ATP: adenosine triphosphate
CHD: coronary heart disease
CPR: cardiopulmonary resuscitation
CVD: cardiovascular disease
ECG: electrocardiogram
IMAC: inner membrane anion channel
MPTP: mitochondrial permeability transition pore
mtDNA-CN: mitochondrial DNA copy number
SCD: sudden cardiac death
SNP: single nucleotide polymorphism

